# A telomere-to-telomere map of somatic mutation burden and functional impact in cancer

**DOI:** 10.1101/2025.10.10.681725

**Authors:** Min-Hwan Sohn, Danilo Dubocanin, Mitchell R Vollger, Youngjun Kwon, Anna Minkina, Katherine M Munson, Samuel FM Hart, Jane E Ranchalis, Nancy L Parmalee, Adriana E Sedeño-Cortés, Jeffrey Ou, Natalie YT Au, Stephanie Bohaczuk, Brianne Carroll, Christian D Frazar, William T Harvey, Kendra Hoekzema, Meng-Fan Huang, Caitlin N Jacques, Dana M Jensen, J Thomas Kolar, Rosa Lee, Jiadong Lin, Kelsey Loy, Taralynn Mack, Yizi Mao, Meranda M Pham, Erica Ryke, Joshua D Smith, Lila Sutherlin, Elliott G Swanson, Jeffrey M Weiss, SMaHT Assembly WG, Claudia Carvalho, Tim HH Coorens, Kelley Harris, Chia-Lin Wei, Evan E Eichler, Nicolas Altemose, James T Bennett, Andrew B Stergachis

**Author notes:** These authors contributed equally to this work (Min-Hwan Sohn, Danilo Dubocanin).

## Abstract

Oncogenesis involves widespread genetic and epigenetic alterations, yet the full spectrum of somatic variation genome-wide remains unresolved. We generated a near-telomere-to-telomere (T2T) diploid assembly of a donor paired with deep short- and long-read sequencing of their melanoma. This revealed that 16% of somatic variants occur in sequences absent from GRCh38, with satellite repeats acting as hotspots for UV-induced damage due to sequence-intrinsic mutability and inefficient repair. Centromere kinetochore domains emerged as focal sites of structural, genetic, and epigenetic variation, leading to remodeling of centromere kinetochore binding domains during tumor evolution. Single-molecule telomere reconstructions uncovered cycles of attrition, deletion, and telomerase-mediated extension that shape cancer telomeres. Finally, diploid chromatin maps exposed that copy number alterations and epimutations, rather than point mutations, predominate in rewiring cancer regulatory programs. These findings define the full landscape of a cancer’s somatic variation and their functional impact, establishing a blueprint for T2T studies of mosaicism.

## Introduction

Cancer genomes are shaped by pervasive genetic and epigenetic alterations that collectively drive oncogenesis, with many of these mutations occurring in the years or decades prior to oncogenesis. Decades of sequencing have revealed recurrent point mutations, copy number alterations, and structural rearrangements across tumor types, yet the full spectrum of somatic variation remains incompletely defined, as our existing maps are grounded in incomplete haploid reference genomes that lack genomic loci that are known to be the most genetically and functionally variable in cancer (*i.e.,* centromeres, telomeres, etc.). Recent advances in telomere-to-telomere (T2T) assemblies and long-read sequencing^1–6^ now permit the comprehensive genetic evaluation of these previously inaccessible genomic regions, and advances in single-molecule chromatin fiber sequencing^7–10^ permit the functional assessment of genetic alterations within these regions.

To comprehensively resolve somatic mutational dynamics and their functional consequences genome-wide, we generated a near-T2T diploid assembly of a donor and paired it with deep short- and long-read sequencing of a melanoma sample obtained from the same donor. This framework enables the unbiased nucleotide-resolution analysis of somatic variants across the entire diploid genome, including sequences absent from GRCh38. Using this framework, we show that satellite repeats serve as genomic sinks for UV-induced mutagenesis, centromere kinetochore domains are hotspots for both genetic and epigenetic variation, and we resolve the relative contribution of telomere erosion and telomerase extension to somatic telomere sequences. Finally, we demonstrate that copy number alterations and epimutations, rather than point mutations, predominantly rewire cancer gene regulatory programs. Together, these findings define the full landscape of somatic variation across a cancer genome, illuminate mechanisms of instability within its most repetitive regions, and provide a blueprint for telomere-to-telomere studies of oncogenesis.

## Results

### Telomere-to-telomere mapping of cancer somatic variants

To establish a genome-wide map of somatic variants in this cancer sample, we first subjected the paired lymphoblastoid cell line (COLO829BL) to PacBio Fiber-seq HiFi sequencing (N50 19.15 kb; 186.24 Gb of sequencing), ultralong Oxford Nanopore sequencing (UL-ONT) (N50 130.85 kb; 186.00 Gb of sequencing), and Hi-C (126.07 Gb of sequencing), enabling us to generate a diploid donor-specific assembly (DSA; 5.92 Gb of genomic sequence with an N50 of 135 Mb) (**Figure 1A** and **1B**, **Table 1** and **Figure S1A** and **S1B**), which we extensively benchmarked for accuracy (estimated QV 62.16 with 98.6% appearing correctly assembled) and completeness of gene content (**Figure S1C**). Using a donor-specific graph genome (DSG) containing the COLO829BL DSA, GRCh38, and CHM13 we identified that the COLO829BL DSA contains 12 chromosomes assembled T2T, with sequence for all 46 human chromosome centromeres, and 188.1 Mb of sequence absent from GRCh38 (3.22% of the COLO829BL genome), establishing the COLO829BL DSA as an accurate and complete representation of the germline genome for the paired melanoma cell line COLO829 that is derived from the same individual as COLO829BL.

**Figure 1.**
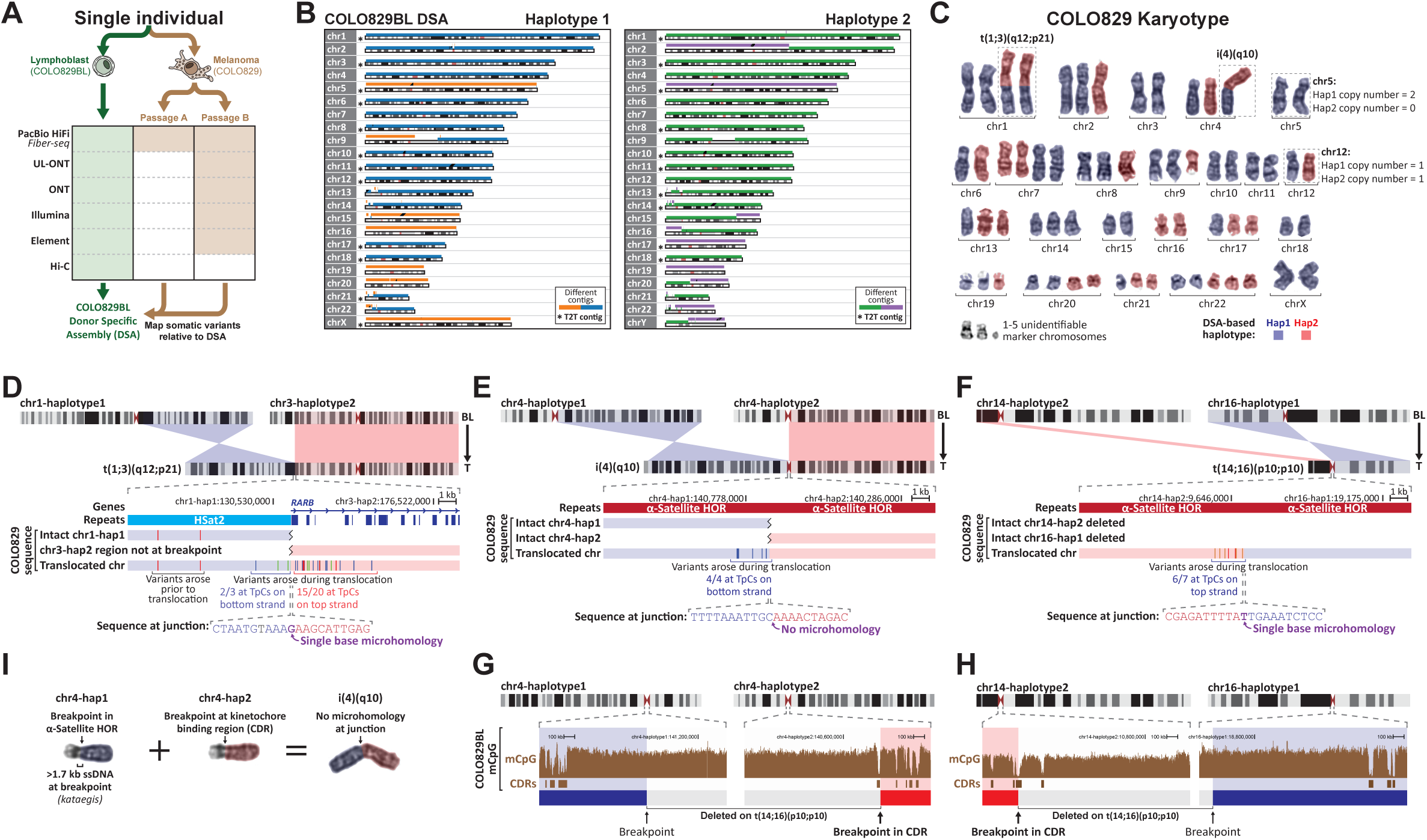
T2T mapping of somatic variants in a melanoma cell line. **A.** Sequencing schematic for a paired lymphoblastoid and melanoma cell line from the same individual, as well as generation of a donor-specific assembly (DSA) from this individual. **B.** Plot of the contiguity of each haploid chromosome in the COLO829BL DSA. **C.** Representative images of each chromosome from the COLO829 melanoma karyotype colored by the presumed haplotype of each chromosome and translocation identity based on read coverage along the COLO829BL DSA. **D.** (top) Ideogram showing the precise breakpoints mediating the t(1;3)(q12;p21) translocation in COLO829 cells based on the COLO829BL DSA. (middle) Sequence differences in COLO829 cells relative to COLO829BL along reads mapping to intact regions of the loci involved in this translocation, as well as the translocated chromosome. (bottom) Sequence at junction with microhomology base in purple. **E.** Same as D, but for the i(4)(p10) chromosome junction. **F.** Same as D, but for a t(14;16)(p10;p10) translocation that was identified using the long-read sequencing data. Note that there are no intact regions of these loci in COLO829 cells. **G.** CpG methylation data in COLO829 cells relative to the breakpoints involved in creation of the COLO829 i(4)(p10). **H.** Same as G, but for the COLO829 t(14;16)(p10;p10) translocation. **I.** Schematic showing the genomic events leading to the formation of the COLO829 i(4)(p10).

**Table 1.** COLO829BL diploid donor-specific assembly and summary statistics.

|  |  | COLO829BL DSA |  |
| --- | --- | --- | --- |
| Assembly materials (Coverage*/N50**) |  | PacBio Fiber-seq (60X/19.15 kb), UL-ONT+ONT (60X/130.85 kb), Hi-C (30X) |  |
| Genome assembly algorithm |  | Verkko v2.1 |  |
| Haplotype |  | Haplotype 1 | Haplotype 2 |
| Number of contigs/scaffolds |  | 56/45 | 58/46 |
| Contig/Scaffold N50 (Mbp) |  | 129.51/147.00 | 130.15/134.93 |
| Number of near T2T contigs/scaffolds |  | 7/9 | 8/11 |
| Sex chromosome |  | chromosome X | chromosome Y |
| BUSCO | Single copy (%) | 98.85 | 95.83 |
|  | Multi copy (%) | 0.69 | 0.67 |
|  | Fragmented (%) | 0.34 | 0.36 |
|  | Missing (%) | 0.12 | 3.14<sup>‡</sup> |
| GC contents (%) |  | 40.8325 | 40.7820 |
| QV |  | 61.6001 | 62.8292 |
| Correctly assembled genome (Gbp/%) |  | 2.962 (98.69) | 2.876 (98.47) |

\* Calculated using 3.1Gb as the denominator.
\*\* If applicable
<sup>‡</sup> The high percentage in this category results from chromosome X genes making up 440 of the 13,780 BUSCO gene sets.

To identify somatic cancer variants within COLO829, we next performed a range of short-read and long-read sequencing on COLO829 cells (**Figure 1A** and **Figure S1A**), using sequencing methods with distinct sequencing error profiles, to accurately assess large-scale chromosomal events, structural variants (SVs), insertions and deletions (indels), and single-nucleotide variants (SNVs) genome-wide. Furthermore, to distinguish cancer somatic variants from those arising during cell culture, we performed PacBio HiFi sequencing on two separate passages of COLO829 (*i.e..* COLO829 passage A, and COLO829 passage B). We then mapped the COLO829 sequencing data to the diploid COLO829BL DSA, and leveraged the DSG to transfer reads and variant calls between the two COLO829BL haplotypes, and GRCh38, enabling us to comprehensively map somatic variants across the entire COLO829BL genome, including within regions absent from GRCh38 in an unbiased manner.

### Kinetochores are hotspots for somatic translocations

Aneuploidies involving whole chromosomes or chromosome arms are a hallmark of many cancers^11,12^, including COLO829 cells^13,14^, which contain whole chromosome duplications and deletions, as well as chromosome arm translocations and isochromosomes (**Figure 1C**). We identified that in COLO829 cells 15 haploid chromosomes have whole chromosome duplications, 6 have whole chromosome deletions, and 6 have chromosome arm deletions/duplications (**Figure 1C**). For example, the four copies of chromosome 20 in COLO829 cells arose from duplications of both haplotypes, whereas the three copies of chromosome 14 arose from a triplication of one haplotype, with the other haplotype being involved in a chromosome arm translocation with 16p.

Emerging data has implicated the kinetochore in the formation of chromosome arm aneuploidies^15,16^, but precise information regarding the relationship between breakpoints and the kinetochore binding region of the centromere has remained elusive owing to the complex genomic architecture of human centromeres^17^. Using the COLO829BL DSA (**Figure S2A**), we precisely delineated the breakpoints underlying all 6 chromosome arm deletions/duplications in COLO829 cells (Figure **S2B-E**). Specifically, we identified that the breakpoints of t(1;3)(q12;p21) localized to a pericentromeric HSat2 array on 1q and the *RARB* gene on 3p (**Figure 1D** and **Figure S2B**). In contrast, all four breakpoints involved in i(4)(q10) and t(14;16)(p10;p10) localized to α-satellite higher order repeats (HOR) (**Figure 1E** and **1F** and **Figure S2C** and **S2D**), regions absent from GRCh38. Two of these three breakpoint junctions had only a single-base microhomology with the third (*i.e.,* i(4)(q10)) having no microhomology. Notably, all three breakpoints were characterized by kataegis^18^ with evidence of 1-5 kb of ssDNA being involved in the breakpoint repair mechanism (**Figure 1D-F**). For example, the t(1;3)(q12;p21) junction was characterized by 20 clustered somatic mutations within 5 kb upstream and 3 clustered somatic mutations 1 kb downstream of the breakpoint, with the far majority (17/23, 73.91%) of these present at TpC dinucleotides, consistent with APOBEC activity on single-stranded DNA (ssDNA). To evaluate for evidence of ectopic recombination between non-allelic homologous regions surrounding these breakpoints, as is seen with break-induced replication (BIR), we focused on t(14;16)(p10;p10), as this chromosome arises from haploid chromosomes that are otherwise deleted in COLO829 cells (**Figure S2E** and **S2F**), enabling us to precisely map the genomic sequence of the translocated chromosome across a >2 Mb region surrounding each breakpoint. This did not reveal any additional somatic structural rearrangements surrounding this junction on t(14;16)(p10;p10). Together, these findings of minimal microhomology with long ssDNA at the sites of these breakpoints suggest that these chromosome arm aneuploidies may result from microhomology-mediated break-induced replication (MMBIR)^19,20^.

To evaluate the relationship between these breakpoints and the kinetochore, we used the COLO829BL long-read CpG methylation data to precisely map the kinetochore binding regions. Specifically, within each chromosome’s α-satellite HOR, the kinetochore is known to localize to specific 20-100 kb regions characterized by CENP-A occupancy, hypo-CpG methylation (*i.e.,* the centromere dip region [CDR])^4^, and dichromatin^9^. Notably, for both i(4)(q10) and t(14;16)(p10;p10), one of the two breakpoints localized to a CDR (Permutation test *p*-value = 0.0048), implicating the kinetochore in mediating these centromere breakages (**Figure 1G** and **1H**). Together, these findings illustrate how mechanisms of cancer somatic alterations within some of the most complex regions of the human genome can be illuminated using a paired DSA approach (**Figure 1I**).

### Genome-wide map of COLO829 sSNVs and their mutational timing

We next sought to extend this analysis by generating a comprehensive genome-wide map of COLO829-specific somatic SNVs (sSNVs). Pairing the COLO829BL DSA and DSG with deep COLO829 long-read PacBio HiFi sequencing, and short-read Element Aviti and Illumina sequencing enabled us to identify and validate SNVs present in both COLO829 passages and absent in COLO829BL (**Figure 2A** and **Figure S3A-C**). Overall, this approach revealed 44,795 sSNVs, and 828 somatic double nucleotide variants (sDNVs), 99.88% of which were validated using UL-ONT sequencing (**Figure S3C**), a technology held out from SNV discovery, indicating a false discovery rate of less than 0.2%. Furthermore, the mutational spectra of these sSNVs and sDNVs match those of UV damage (**Figure 2C**), consistent with these variants arising during formation of the melanoma, as opposed to being culture artifacts^21,22^. Overall, 15.6% of these sSNVs mapped to regions of the COLO829BL DSA absent from GRCh38 (**Figure 2A**), including 599 sSNVs that map to non-repetitive loci that are uniquely present within the DSA, indicating that GRCh38-based somatic variant catalogs systematically underrepresent the true extent of somatic variation that exists in a sample.

**Figure 2.**
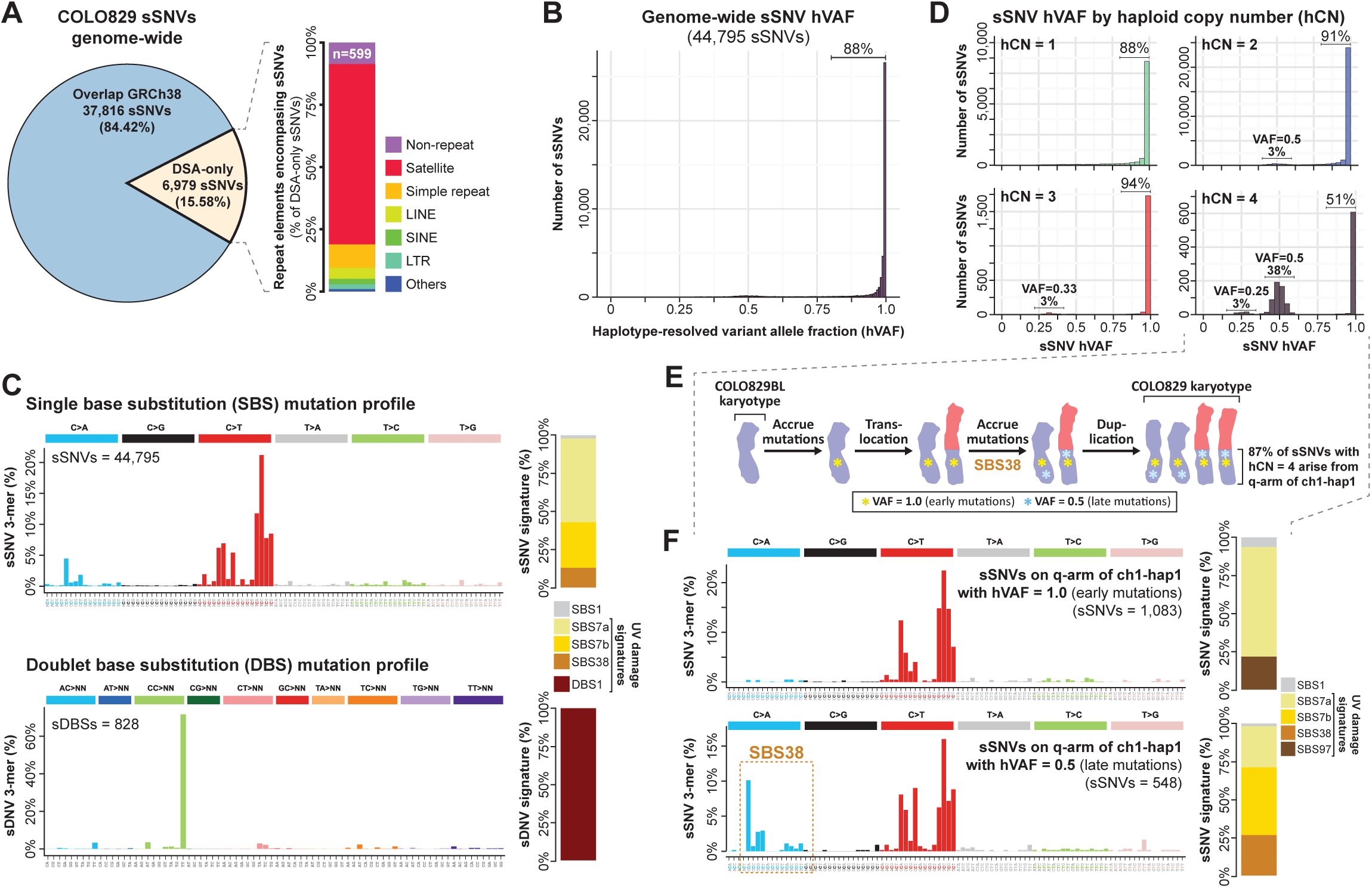
Genome-wide mapping of somatic SNVs and their timing in COLO829. **A.** Pie chart showing the proportion of COLO829 sSNVs that mapped to regions with paired sequence in GRCh38 reference. Inset showing stacked bar plot of the repeat annotations for variants that are present in genomic regions unique to the COLO829BL DSA. **B.** Haploid variant allele fraction (hVAF) for all COLO829 sSNVs along the COLO829BL DSA. **C.** Frequency of all sSNVs and sDBSs within different k-mer contexts genome-wide (left), as well as contribution of known COSMIC signatures on each of the mutational spectra (right). **D.** Distribution of hVAF separated by different somatic haploid copy number (hCN) states. **E.** Schematic history of mutational events that involve chromosome 1 and 3, inferred by hVAF and hCN across the corresponding chromosomes as well as long-reads spanning the junction. **F.** SBS mutational spectra within the q-arm of the chromosome 1 haplotype involved in the translocation event, separated by early and late events by hVAF range, and the contribution of known COSMIC UV-associated signatures, respectively.

Although the far majority of sSNVs have a variant allele fraction (VAF) consistent with being fixed within one of the two haplotypes (**Figure 2B**), 38% of sSNVs along the q-arm of chromosome 1 haplotype 1 (chr1q_hap1_), which has a haploid copy number of 4 (hCN4), showed a VAF of ∼0.5 (**Figure 2D**). Chr1q_hap1_ is present in 2 copies from a duplication of the intact chromosome 1, and 2 copies from a duplication of t(1;3)(q12;p21) (**Figure 1C**), and long-read sequencing data within the chr1q_hap1_ HSat2 array revealed that encompassed sSNVs with a haploid VAF of 0.5 are exclusively present on either t(1;3)(q12;p21) or on the intact chromosome 1 (**Figure S3D** and **S3E)** implicating t(1;3)(q12;p21) as the first retained chromosome gain aneuploidy in the lifespan of the cell that gave rise to COLO829 (**Figure 2E**). Notably, although chr1q_hap1_ sSNVs with a VAF of 1.0 and 0.5 both had mutational spectra consistent with arising from UV damage, sSNVs with a VAF of 0.5 (*i.e.,* late clonal variants) uniquely contained C>A transversions that matched the mutational signature SBS38 (**Figure 2F**), providing further evidence that SBS38 is uniquely present within the later stages of melanoma evolution ^23^. Together, these findings establish the validity of our genome-wide DSA-based assessment of cancer somatic variants, including in genomic regions absent from the GRCh38 reference.

### Satellite repeats act as genomic sinks for UV damage sSNVs

As our map of sSNVs comprehensively captures somatic variants genome-wide, including within complex loci absent from GRCh38 (**Figure 2A**), it enables us to accurately quantify the relative rate of somatic variants and their mutational mechanisms across different genomic loci. Comparing the rate of sSNVs across different repeat elements demonstrated that overall repeat elements harbor an elevated rate of sSNVs within COLO829 (1.4-fold higher than non-repeat elements), with satellite elements harboring the highest rate of sSNVs (3.79-fold higher than non-repeat elements) (**Figure 3A**). Furthermore, we observed a 5.05-fold difference in the rate of sSNVs among different classes of satellite repeats, with HSat2 elements showing the highest rate of sSNVs at 6.1×10^-5^ SNVs/bp (7.78-fold higher than non-repeat elements). Notably, HSat2 and other satellite repeats showed a consistently elevated rate of sSNVs across these extended repeats (**Figure 3B**), suggesting that an intrinsic property of these repeats may be underlying their elevated mutation rate.

**Figure 3.**
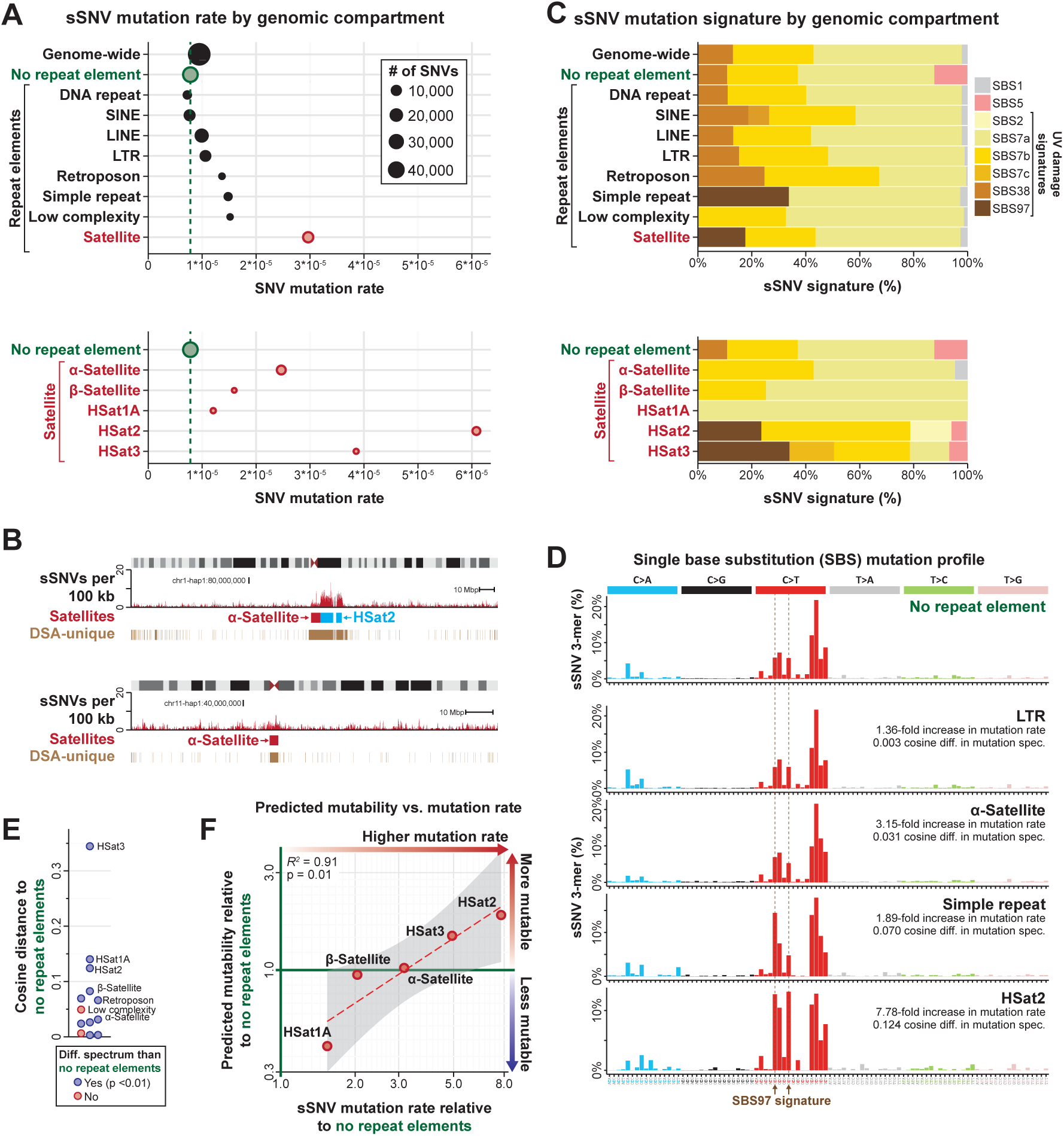
Satellite repeats act as genomic sinks for UV damage sSNVs. **A.** Rate of sSNVs across various repeat element classes along the COLO829BL DSA. (Bottom) Rates of sSNVs across various satellite repeats. **B.** Density of sSNVs within 100kb bins across the entire chromosome 1-haplotype1 and chromosome 11-haplotype 1 in COLO829 cells. Alpha-satellites, HSat2 arrays, and regions uniquely present in the COLO829BL DSA indicated. **C.** Contribution of known UV-associated COSMIC signatures to the 3-mer mutational spectra within various repeat element classes along COLO829BL DSA. **D.** Frequency of sSNVs within different k-mer contexts across different genomic regions. Indicated to the right is the difference in the mutation rate and mutational spectrum of that genomic region relative to the non-repeat regions of the COLO829BL DSA. **E.** Swarm plot of cosine distance of the mutational spectrum of a genomic region relative to the non-repeat regions of the COLO829BL DSA. Genomic regions with a significant difference in their mutational spectra are indicated by color. **F.** Scatter plot showing the relationship between the predicted mutability of a given satellite repeat relative to non-repeat elements along the COLO829BL DSA and the observed sSNV mutation rate of a given satellite repeat relative to non-repeat elements along the COLO829BL DSA. Confidence interval for linear fit shown in grey.

To evaluate the mechanism(s) driving satellite elements to have markedly elevated mutation rates, we first compared the *k*-mer normalized mutational spectra of different repeat classes (**Figure 3C** and **Figure S5A**). This revealed that all of the repeat classes, including the different satellite repeats, largely have similar normalized mutational spectra, which are dominated by UV-specific signatures (**Figure 3C-E**). This indicates that similar mutational processes underlie sSNVs in both repetitive and non-repetitive genomic loci in COLO829 (**Figure S5B**). To further resolve what could be driving the marked differences in the rate of sSNVs between the different satellite repeat classes, which should all be late replicating^24^, we quantified the predicted mutability of each satellite class based on its enrichment of *k*-mers that preferentially harbor sSNVs in non-repeat elements within COLO829 cells (**Figure 3F** and **Figure S5C**). This revealed that the mutation rate within satellite repeats was significantly associated with the predicted mutability of the underlying sequence (*R^2^* = 0.91, *p*-value = 0.01) (**Figure 3F**), indicating that satellite repeats with the highest sSNV rate are preferentially populated by *k*-mers that are exquisitely sensitive to UV damage. However, even after controlling for predicted mutability, satellites still have a markedly elevated mutation rate compared to non-repeat elements (**Figure 3F**), suggesting that either satellites are more prone to UV damage or that the repair machinery for resolving UV damage is less efficient in satellites. Together, these findings indicate that satellite repeats, which comprise 4.27% of this donor’s genome, are exquisitely prone to accrue UV mutations.

### Focal kinetochore occupancy is marked by both somatic genetic and epigenetic variation

Given the propensity for chromosome arm translocation breakpoints to localize to the kinetochore binding region of the centromere, we next wanted to evaluate whether the kinetochore binding region showed altered rates of somatic SNVs. The COLO829BL DSA contains complete maps of the CDRs for 43 out of the 46 haploid chromosomes (93.48%), enabling us to evaluate mutational processes that may be uniquely occurring within these regions. Overall, we observed that COLO829BL CDRs contain a significantly higher rate of COLO829 SNVs relative to the rest of the α-satellite HORs (4.20×10^-5^ vs 2.32×10^-5^, respectively, Two-proportion Z-test *p*-value = 3.05×10^-31^) (**Figure 4A** and **4B** and **Figure S6A**). However, the mutational pattern within CDRs largely mirrors the rest of the α-satellite HORs, which is predominantly driven by UV damage signatures (**Figure 4B**), and CDRs contain a similar *k*-mer profile as the rest of the α-satellite HORs (**Figure S6B** and **S6C**), indicating that the higher rate of mutations in CDRs is not reflective of a distinct mutational process, but rather either reflects a higher rate of UV-induced DNA lesions, or a depressed ability to repair these lesions.

**Figure 4.**
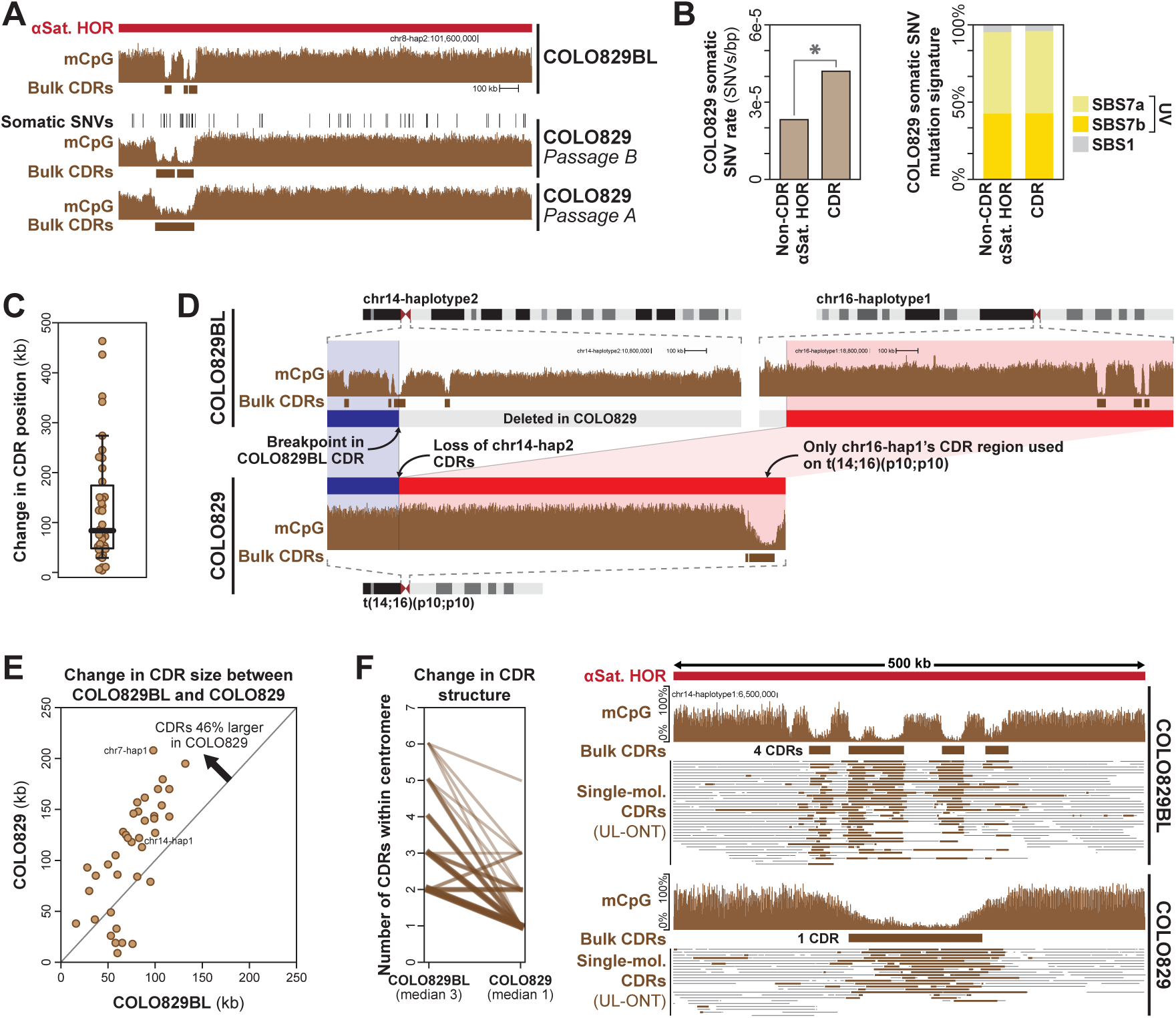
Somatic genetic and epigenetic variation within centromeres. **A.** Genomic locus showing CpG methylation data from COLO829BL and two passages of COLO829 within the chromosome 8-haplotype 2 centromere alongside the centromere dip regions (CDRs) that correspond to kinetochore binding regions. Also shown are COLO829 sSNVs. **B.** (left) Rate of COLO829 sSNVs within CDR and non-CDR regions of COLO829BL alpha-satellite HORs (*, *p*-value <0.01). (right) Contribution of known UV-associated COSMIC signatures to the 3-mer mutational spectra within CDR and non-CDR regions of COLO829BL alpha-satellite HORs. **C.** Average change in the CDR location between COLO829 and COLO829BL cells. **D.** Genomic locus showing CpG methylation data from COLO829BL and COLO829 along the intact copies of chromosome 14 and 16 in COLO829 cells, as well as the translocated copy of t(14;16)(p10;p10) in COLO829 cells. **E.** Scatter plot showing the overall size of the CDRs for each centromere in COLO829BL and COLO829. Shown is the average difference above. **F.** (left) For each centromere, shown is the number of CDRs within that centromere in COLO829BL versus COLO829 cells, as well as the median number below. (right) genomic locus showing per-molecule CDR calls using the UL-ONT data from both COLO829BL and COLO829 showing the condensation of the CDRs from COLO829BL into one large CDR in COLO829 for chromosome 14-haplotype 1.

We next wanted to determine whether CDRs were also hotspots for epigenetic variation in addition to genetic variation. Prior work has identified that CDR positions are largely stable from parent to offspring^6^. However, CDRs harboring *de novo* structural variants have been observed to shift their location by ∼260LJkb^6^. Overall, we observed that 45% of COLO829 CDRs shifted their center of mass by >100 kb relative to COLO829BL (**Figure 4C**). Furthermore, along t(14;16)(p10;p10), the native chr14-hap1 CDR was silenced in COLO829 cells, and instead only a single CDR located 1.7 Mb away was utilized in COLO829 cells (**Figure 4D**), indicating the ability of α-satellite HORs to accommodate kinetochore binding regions at multiple locations. In addition to shifting their location, we also observed large-scale reorganization of the size and structure of CDRs between COLO829BL and COLO829. Specifically, whereas each COLO829BL centromere was marked by ∼3 discrete CDRs containing a combined total of ∼52 kb of sequence, COLO829 centromeres were marked by only ∼1 CDR containing a combined total of ∼94 kb of sequence (**Figure 4E** and **4F**). Notably, these new CDR locations appeared to be mitotically stable, as they were stable across both passages of COLO829 cells (**Figure 4A**), and the single-molecule UL-ONT mCpG data showed consistent single-molecule CDR patterns in COLO829 cells (**Figure 4F**). Together, these findings indicate that the kinetochore binding regions of centromeres are hotbeds for both somatic genetic and epigenetic variation during COLO829 oncogenesis.

### Mechanistic basis for oncogenic alterations in telomere length

Oncogenesis is known to be associated with pervasive changes in telomeres. Whereas some cancers utilize the alternative lengthening of telomeres (ALT) pathway to maintain telomeres, many cancers are known to acquire somatic mutations in the *TERT* promoter that result in increased *TERT* expression and telomerase activity^25^. COLO829 contains a chr5:1295113_4GG>AA DNV within the promoter of *TERT*, creating a *de novo* E26 transformation-specific (ETS) transcription factor binding motif that has been previously described to be critical for enhancing *TERT* promoter activity in various cancers^26–28^. However, despite the presence of this activating *TERT* mutation in COLO829 cells, our long-read sequencing data revealed that the COLO829 telomeres were on average 2.3 kb shorter than those in COLO829BL (*p*-value = 2.49×10^-78^, Mann-Whitney U) (**Figure 5A** and **5B**). To investigate the relative contribution of telomere attrition and *de novo* elongation, we developed an approach that leverages encompassed telomere variant sequences (TVRs) to resolve the historical state of each telomere molecule. TVRs are non-TTAGGG repeats that are frequently found within the proximal ends of most telomeres^29–31^, and prior work has shown that TVRs can be somatically restructured into canonical TTAGGG repeats via cycles of telomere shortening and subsequent *de novo* telomerase elongation^30^.

**Figure 5.**
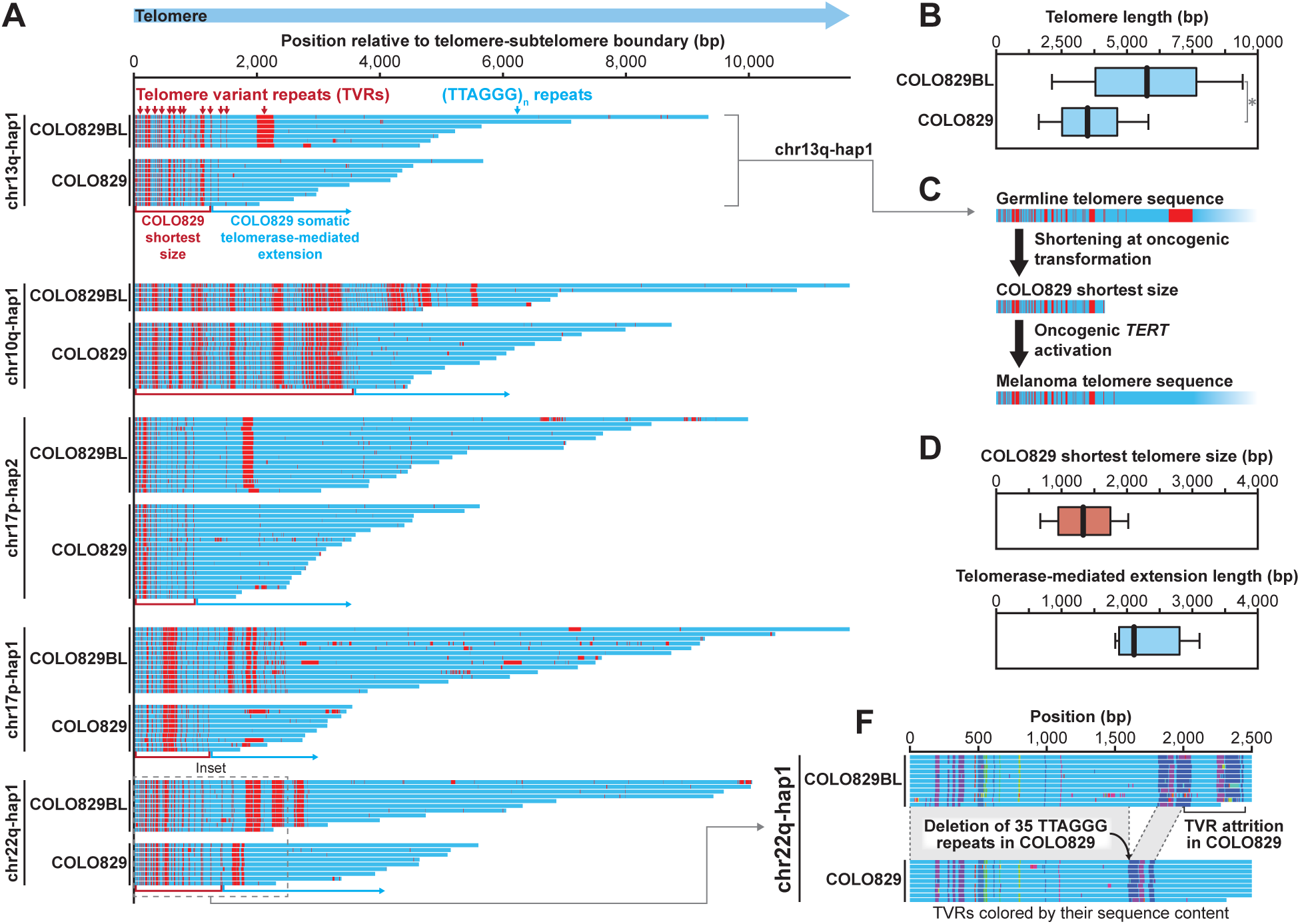
Somatic telomere alterations in COLO829 cells. **A.** Single-molecule ONT genetic architectures of individual telomeres in COLO829BL and COLO829 cells with the subtelomere-telomere junction at the left side, and the individual telomere molecules colored based on whether the sequence contains a canonical telomere TTAGGG sequence or a TVR. **B.** Box-and-whisker plot showing the distribution of single-molecule telomere length measurements in COLO829BL and COLO829 cells based on ONT sequencing data. **C.** Schematic showing the process of telomere attrition followed by telomerase-mediated extension during oncogenic transformation of COLO829 cells based on the TVR fingerprints along the chr15q-hap1 telomere. **D.** Box-and-whister plots showing the (top) lengths of the inferred shortest telomere size and (bottom) lengths of the somatic telomerase-mediated extension of each measurable telomere based on comparison of the pattern of TVRs at that telomere in COLO829BL and COLO829 cells. **E.** Single-molecule ONT genetic architectures of the chr22q-hap1 telomere in COLO829BL and COLO829 cells with the subtelomere-telomere junction at the left and each TVR colored based on its underlying genetic sequence displaying the (TTAGGG)_∼35_ deletion between TVRs.

Comparing the per-molecule telomere sequencing data from COLO829BL and COLO829 revealed that the base of every telomere had a similar TVR pattern between these two samples (**Figure 5A**). However, TVRs on 15.6% (n=64, 12 (18.1%) with quantitatively detected restructuring, 2 filtered out visually to 10 total) of telomeres had distinct TVR patterns between these two samples (**Figure 5A**), characterized by TVRs that are present in the COLO829BL sample being largely replaced with canonical TTAGGG repeats in COLO829 (**Figure 5A**), consistent with a model whereby this pattern reflects telomere attrition prior to oncogenesis followed by *de novo* telomerase-mediated extension following the acquisition of the promoter *TERT* variant. Using these TVRs as single-molecule genomic markers, we calculated that telomeres shortened on average to within 1.4 kb of the telomere/subtelomeric junction prior to the acquisition of the promoter *TERT* variant, which was followed by approximately ∼2.3 kb of somatic telomerase-mediated extension (**Figure 5C** and **5D**). Notably, these TVRs also exposed evidence of somatic deletions internal to the telomere, as can be seen in the COLO829 chr22q-hap1 telomere (**Figure 5E**). Together, these findings enable us to reconstruct the length and sequence of each telomere at the time of oncogenesis, revealing the relative contribution of erosion, somatic structural variants, and *de novo* telomerase-mediated extension to cancer telomere sequence and length. This highlights that although the COLO829 telomeres are significantly shorter than those from COLO829BL, this does not represent a continual erosion of the germline telomeres, but rather represents a single erosion event followed by subsequent telomerase-mediated extension and then steady-state telomere maintenance.

### Somatic CNVs and epimutations predominantly rewire the gene regulatory landscape of COLO829

We next sought to resolve the impact of the identified COLO829 somatic variants on the gene regulatory program of COLO829 cells by leveraging the paired haplotype-phased Fiber-seq data from both COLO829BL and COLO829 (**Figure S9A** and **S9B**). To quantify whether somatic variants within COLO829 are associated with altered chromatin, we leveraged the fact that an individual somatic variant will only be present on one haplotype within COLO829, so if both haplotypes are retained in COLO829, we can use the opposite haplotype as a control, as the opposite haplotype will be subjected to the same trans environment as the one containing the somatic variant. Of the 29,189 COLO829 sSNVs and sDNVs that localize to loci that retain both haplotypes, only 280 overlap a regulatory element that is accessible on at least one of the two haplotypes in COLO829 cells, consistent with regulatory DNA having an overall reduced cancer mutation rate (*p*-value = 1.091×10^−14^, Pearson’s Chi-squared test with Yates’ continuity correction)^32^. However, only 3.93% (11/280) of these somatic variants were associated with a significant change in chromatin accessibility on the haplotype that contains the somatic variant, with 9 showing a decrease in chromatin accessibility, and 2 showing an increase in chromatin accessibility (**Figure 6A** and **Figure S9C**). For example, one of these 11 sSNV appears to ablate chromatin accessibility and TF occupancy at a CTCF binding element located within a segmentally duplicated regulatory element with >99% sequence identity at two locations along chromosome 2 (**Figure 6B**), highlighting the power of our approach for detecting functional sSNVs within genomic loci traditionally marred by mapping artifacts.

**Figure 6.**
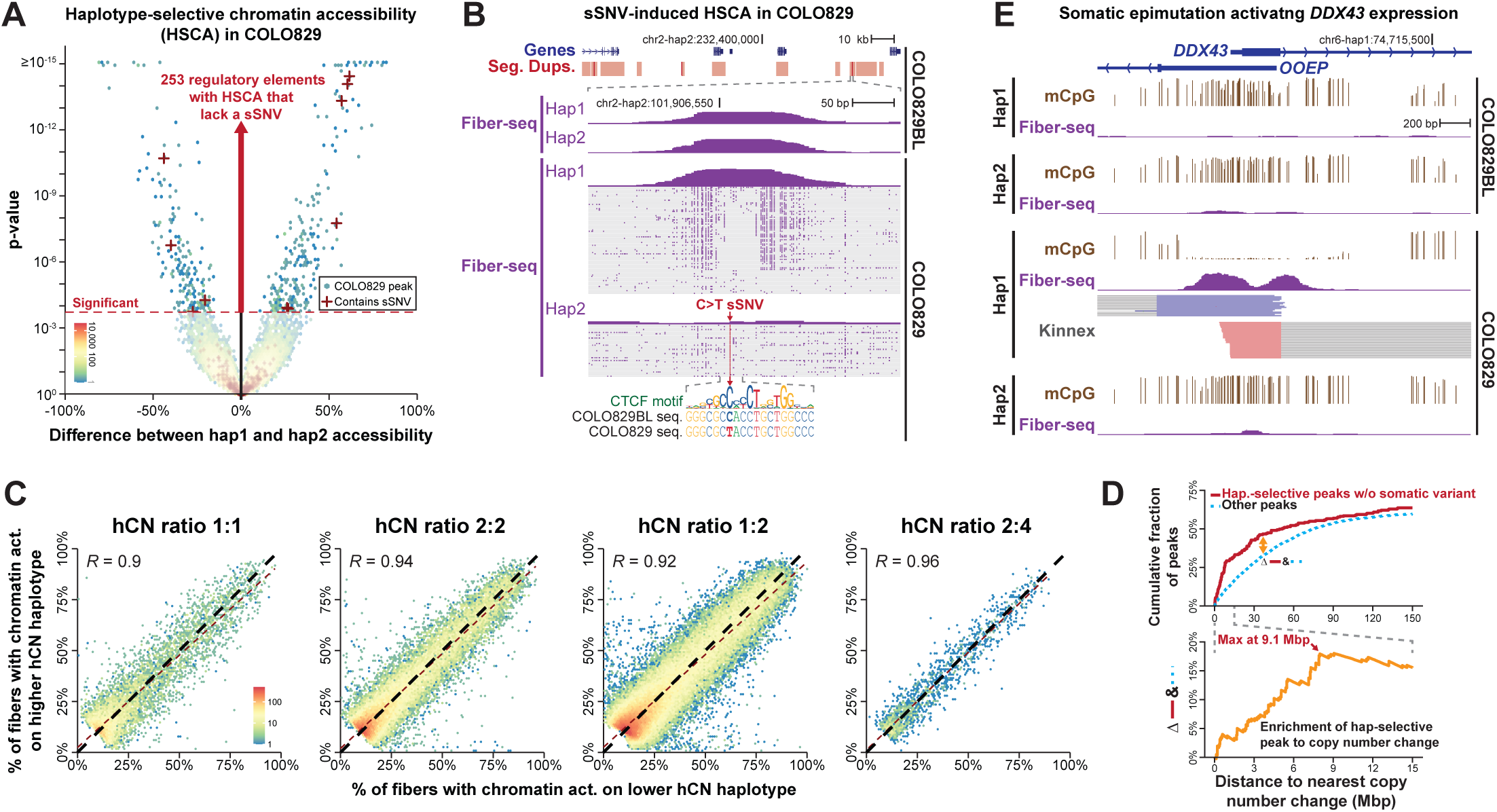
The impact of somatic variants on the epigenome. **A.** Volcano hexagonal binned plot showing the relative Fiber-seq chromatin accessibility at all Fiber-seq peaks in COLO829 cells that are not differentially accessible in COLO829BL cells. This is limited to peaks contained within genomic loci where both haplotypes are retained in COLO829 cells. Significance determined using Fisher’s exact test with Benjamini-Hochberg multiple testing correction (FDR < 0.05). Red crosses mark elements with haplotype-selective chromatin accessibility and overlapping COLO829 sSNVs. **B.** Pseudobulked and single-molecule Fiber-seq chromatin data from COLO829BL and COLO829 cells showing a COLO829 sSNV located within a segmentally duplicated region that disrupts protein occupancy at a CTCF binding site selectively on haplotype two of COLO829. **C.** Hexagonal binned plots showing for each regulatory element along the COLO829 genome the percentage of chromatin fibers that have an actuated regulatory element at that site by haplotype based on COLO829 Fiber-seq data. Four plots are separately displaying Fiber-seq elements that are present within genomic loci with different haploid copy number configurations. **D.** (top) The fraction of haplotype-selective Fiber-seq peaks without somatic variants or heterozygous germline variants (red) or other Fiber-seq peaks (dashed blue) located within a specific distance (Mbp) of a change in copy number along the COLO829 genome. (bottom) The difference in red and dashed blue line above. **E.** Pseudobulked Fiber-seq chromatin and CpG methylation data and long-read transcript data from COLO829BL and COLO829 cells showing a promoter that is selectively activated on haplotype 1 in COLO829 cells via a somatic epimutation that creates chromatin accessibility and hypo-CpG methylation.

However, COLO829 cells contained an additional 253 regulatory elements with haplotype-selective chromatin accessibility that could not be explained by somatic genetic variants, imprinted elements, or functional germline heterozygous variants (**Figure 6A** and **Figure S9D**), suggesting that additional genetic or epigenetic features may be playing a dominant role in modulating the chromatin epigenome of COLO829 cells. As 78.05% of the COLO829 genome contains a somatic CNV (sCNV) relative to COLO829BL, we next sought to investigate the role of sCNVs in altering the haplotype-selective chromatin epigenome of COLO829 cells. sCNVs have the potential to alter the chromatin epigenome via additive or non-additive effects. For example, if both copies of a duplicated genomic locus have identical function to that of the non-duplicated copy, the sCNV would be considered to act in an additive manner as there would be parity between the dosage of the encompassed regulatory elements and their functional output. In contrast, if one of the duplicated copies has altered function (*e.g.,* from positional effects of the sCNV, or from somatic epimutations), then the sCNV would be considered to act in a non-additive manner as there would not be parity between the genetic dosage of the encompassed regulatory elements and their functional output. Overall, we observed that COLO829 sCNVs systematically have an additive impact on the chromatin epigenome, with copy number gains resulting in a synchronous increase in chromatin accessibility (Pearson’s *R* = 0.92, *p*-value < 2×10^-16^) (**Figure 6C** and **Figure S9E**). However, these 253 elements appeared to have a non-additive effect, raising the possibility that their chromatin accessibility is being impacted by positional effects.

To quantify the impact of positional effects on these 253 elements with non-additive chromatin accessibility, we assessed their localization relative to sCNV boundaries in COLO829 cells, revealing that these 253 elements were significantly enriched for being adjacent to CNV boundaries (**Figure 6D**). However, this enrichment extended up to 9 Mb away from the sCNV breakpoints (**Figure 6D** and **Figure S9F**), well beyond the typical length of chromatin loops or topologically associating domains (TADs), implying that these non-additive effects are caused by the emergence of somatic epimutations^10^ arising along duplicated regulatory programs. For example, *DDX43*, which is a gene involved in melanoma oncogenesis^33^, appears activated in COLO829 cells via a somatic epimutation that selectively derepresses its promoter along haplotype 1 (**Figure 6E**). Together these findings indicate that additive effects of sCNVs, and somatic epimutations play a dominant role in rewiring the chromatin epigenome of COLO829, with only 0.02% of sSNVs having an observable direct impact on the chromatin epigenome.

## Discussion

Here, we establish a roadmap for generating comprehensive, unbiased maps of somatic variation across an entire diploid genome and illuminate how somatic genetic and epigenetic alterations within repetitive elements shape genome structure and function during oncogenesis. By integrating long-read sequencing, diploid assembly, single-molecule chromatin fiber sequencing, and graph genomes, we overcome the limitations of incomplete references such as GRCh38, CHM13, and reference pangenomes. This framework enables somatic variants to be defined directly within the context of an individual’s genome, providing a new standard for somatic mutation characterization in precision cancer genomics.

Our analysis identifies satellite repeats, regions systematically absent from GRCh38-based catalogs, as major hotspots of somatic mutagenesis. Orthogonal validation across sequencing platforms, variant allele fraction analyses, and mutational signature profiling confirmed the accuracy of these excess mutation calls. After correcting for the mutability of encompassed *k*-mers, as measured in non-repetitive regions, satellites displayed a threefold higher mutation rate than non-repeats, far exceeding the elevation seen in late-replicating euchromatin. This work extends upon recent studies that show that germline^6^ and early post-zygotic^34^ mutations preferentially localize to repeat elements. However, these germline and early post-zygotic mutations are largely driven by an elevated rate of clock-like mutations and gene conversion events^6,34^, which is in contrast to the largely UV-driven processes seen in COLO829. Specifically, although the different satellite subclasses in COLO829 displayed marked differences in their mutation rates, these differences are largely explained by their relative abundance of UV-damage-sensitive *k*-mers, indicating that a satellite region’s sensitivity to UV-damage and/or its ability to correctly repair UV-damage is playing a dominant role in modulating its sSNV mutation rate. These findings point to elevated incidence and/or impaired repair of UV lesions^35^, rather than a satellite-specific mutagenic process, as the driver of their elevated somatic mutation burden.

Centromeres, and particularly kinetochore binding domains, represent an extreme example. These regions exhibit a markedly elevated sSNV rate even after adjusting for sequence context, and they serve as recurrent sites of chromosome arm translocations. Notably, breakpoints within α-satellite HORs are characterized by clustered mutations with a pattern matching that of kataegis. Kataegis has been shown to result from break-induced replication (BIR) repair in yeast models^36^, wherein a long 5’-3’ DNA resection is targeted by APOBEC cytidine deaminases. However, neither of the two breakpoint junctions we observed within α-satellite HORs contained apparent homology arms, with one having a single-base microhomology, and the other having no homology at all. Together, these data implicate replication stress and deficient repair within kinetochore binding domains as a mechanistic basis for the prevalence of centromeric breakpoints and aneuploidies in cancer. Notably, it is likely that these findings are not unique to COLO829 or UV-damage, as germline and early post-zygotic mutation rates have similarly been found to be elevated in satellites^34^ and the kinetochore binding region in particular^6^, mutational processes that are not UV-driven.

Telomere reconstructions further reveal that oncogenic transformation was associated with a single bottleneck of extensive attrition, followed by stochastic telomerase-mediated extension. This punctuated model contrasts with gradual erosion and provides a molecular record of the minimal telomere length tolerated prior to *TERT* reactivation. Extending this approach to additional paired tumor-normal genomes should clarify the thresholds of telomere dysfunction that precede immortalization.

Finally, we identify pervasive somatic epimutations throughout the COLO829 genome. Specifically, nearly half of centromeres underwent large-scale repositioning of kinetochore binding regions, in some cases by more than a megabase, highlighting the remarkable plasticity of α-satellite sequences in accommodating stable alternative centromere locations. In addition, the epigenetic size of a centromere’s CDR systematically expanded in COLO829 cells, consistent with microscopy-based measurements in transformed cell lines^37^. Genome-wide, copy number alterations acted largely additively on chromatin accessibility^38^, whereas epimutations^10,39^ accounted for non-additive effects, including activation of known oncogenes such as *DDX43*. Strikingly, only 0.02% of somatic SNVs directly altered regulatory chromatin, underscoring that structural and epigenetic variation, rather than point mutations, play the predominant role in reshaping cancer gene regulation.

In sum, this study delivers the first near–telomere-to-telomere map of somatic variation in cancer, revealing how repetitive DNA, centromeres, telomeres, and epimutations collectively sculpt the cancer genome. Beyond defining the mutational burden of a single tumor, our framework establishes a blueprint for future telomere-to-telomere studies of oncogenesis and genomic mosaicism.

## Methods

### Sequencing data generation

All COLO829BL and COLO829 sequencing data used in this study were generated as part of the Somatic Mosaicism across Human Tissue (SMaHT) benchmarking^40^ study, "Comprehensive benchmarking of somatic mutation detection by the SMaHT Network." The SMaHT consortium established COLO829BL/COLO829 as one of the reference standards for systematically evaluating somatic variant detection. The dataset we leveraged encompasses comprehensive genomic and epigenomic profiling across multiple sequencing platforms and technologies including ultra-long Oxford Nanopore Technology (ONT) sequencing, standard ONT sequencing, standard PacBio HiFi sequencing, single-molecule PacBio HiFi chromatin fiber sequencing (fiber-seq) for chromatin accessibility profiling, and short-read whole-genome sequencing using Illumina and Element AVITI Biosciences platforms. Detailed information regarding COLO829BL/COLO829 cell culture, library preparation and sequencing protocols can be found in the flagship paper^40^. All the data can be accessed from the SMaHT Data Portal at https://data.smaht.org/.

### Building COLO829BL diploid genome assembly

To construct the COLO829BL donor-specific assembly, we used 60x coverage of PacBio HiFi and Oxford Nanopore Technologies (ONT) data, along with 30x coverage of Hi-C data. To maximize N50 of the assembly, ONT reads were downsampled by selecting the longest reads until a targeted 60x coverage was achieved. Specifically, based on the index file (.fai) of the ONT FASTQ, read IDs were extracted in descending order of read length until reaching 60× coverage based on 3.1 Gbp as the genome size. Reads corresponding to these IDs were then stored separately to generate the downsampled ONT FASTQ file. Phased assembly was generated using Verkko v2.1^2^. After the initial assembly, we implemented additional filtering steps to ensure that contaminated sequences are removed using NCBI FCS^41^ v0.4.0 beta and BLAST^42^ v2.15. In short, contigs shorter than 10 bp, which can cause errors in FCS, were excluded, followed by the detection of foreign DNA and adapter sequences. Subsequently, mitochondrial, Epstein–Barr virus (EBV) and ribosomal DNA (rDNA) sequences were identified using a BLAST-based approach. The detected contaminant and adapter regions were trimmed, and rDNA contigs were separated from the assembly. The command used to detect foreign DNA is as follows: *python3 fcs.py --image FCS_IMG screen genome --fasta HAPLOID_ASSEMBLY.fasta --out-dir OUTDIR --gx-db GXDB_LOC --tax-id 9606*. The command used to detect adapter sequence is as follows: *av_screen_x -o OUTDIR --euk HAPLOID_ASSEMBLY.fasta*. The command used to detect mitochondrial, EBV, and rDNA sequences is as follows: *blastn -query HAPLOID_ASSEMBLY.fasta -db TARGET_DB -outfmt 6 –evalue 1e-30 > OUTPUT*. Finally, for each haplotype, a decontaminated assembly FASTA file was produced with foreign DNA (including EBV), adapter, mitochondrial sequences, and rDNA contigs removed.

### Evaluating quality of COLO829BL DSA

Comprehensive evaluation of the COLO829BL DSA was essential as all subsequent analyses would directly depend on its accuracy and completeness. Therefore, we utilized multiple orthogonal assessments and different metrics to validate the quality of the assembly.

We first assessed the contiguity of the DSA from N50 and NG50 metrics (based on 3.1Gbp genome size) using calN50 (https://github.com/lh3/calN50). Then, we calculated QV scores using the combination of Meryl^43^ v1.4.1 (*meryl k=21 count ILLUMINA_READ.fastq.gz output READS_DB.meryl*) and Merqury^43^ v1.3 (*merqury.sh READS_DB.meryl HAP1_ASSEMBLY.fasta HAP2_ASSEMBLY.fasta MERQURY_OUTPUT*) to estimate the sequence accuracy of the COLO829BL assembly. QV values were derived as phred-scaled scores for each haplotype, based on error rates estimated by comparing the k-mer database generated from Illumina paired-end sequencing data of COLO829BL with the assembly sequences.

Next, we assessed the gene completeness of the assembled haplotypes using compleasm^44^ v0.2.6, based on the presence and integrity of 13,780 known single-copy orthologs included in the primates_odb10 database (*compleasm run --mode busco -L DB_DIR -l primates_odb10 -o OUT_DIR -a HAPLOID_ASSEMBLY.fasta*).

Furthermore, we implemented genome assembly evaluation tools to systematically identify problematic genomic regions within the DSA, and these areas are removed from all subsequent analyses, unless stated otherwise (hereby referred to as “potentially misassembled regions”). Specifically, regions identified as Erroneous, Duplicated, or Collapsed by Flagger^5^ v0.3.3 as well as regions flagged as MISJOIN, COLLAPSE, COLLAPSE_VAR, COLLAPSE_OTHER or HET, by NucFlag (https://github.com/logsdon-lab/NucFlag), a wrapper for NucFreq^45,46^ framework, are excluded. These quality control steps ensure a series of reliable computational analyses against the DSA, preventing assembly artifacts from introducing potential bias or errors into subsequent biological interpretations or statistical analyses. For Flagger, PacBio HiFi reads were aligned to the combined haplotype 1 and haplotype 2 diploid assembly using minimap2^47,48^ v2.28 (*-Y -x map-hifi --eqx -L --cs*), after which the Flagger WDL pipeline was executed (*java -jar cromwell.jar run flagger_end_to_end.wdl --inputs INPUT.json --metadata-output OUTPUT.json*). For NucFlag we leveraged alignment from COLO829BL PacBio Fiber-seq data aligned onto the DSA with an added filtering step to retain only the primary alignment reads (see “Alignment and post-processing of reads” section). NucFlag was run with the default configuration.

To determine the chromosome of origin for contigs in the COLO829BL DSA, we mapped the assembly to the T2T-CHM13^1^ reference genome using the Snakemake workflow designed to handle assembly-to-assembly alignment (Data and code availability). Briefly, DSA contigs are aligned onto T2T-CHM13 using minimap2^47,48^ with the following parameter: *-x asm20 --eqx --cs*

*--secondary=no*. Then, we implemented rustybam (10.5281/zenodo.15580393) *trim-paf* command (*--remove-contained*) to prevent aligning the same section of DSA contigs to multiple stretches of the target sequence. Using the resulting pairwise alignment records, we calculated for each DSA contig the percentage of its bases aligned to each T2T-CHM13 chromosome by summing all alignment block lengths to each chromosome and dividing by the total contig length. The chromosome of origin for each DSA contig was assigned as the T2T-CHM13 chromosome with the highest alignment percentage.

A contig was classified as telomere-to-telomere (T2T) if it contained telomeric sequences ≥100 bp on both ends. Telomeric sequences were determined by seqtk (https://github.com/lh3/seqtk) *telo* command. Furthermore, to locate the centromeres in the DSA, we identified the coordinates in the DSA that correspond to the centromere annotation in T2T-CHM13 with rustybam *liftover* command by using the pair-wise alignment information between the two assemblies created to identify chromosome of origin above. The leftmost and rightmost coordinates from the output for each contig is used to establish approximate centromere locations. Within these regions, we identified "ALR_Alpha" satellite repeats (annotated by RepeatMasker below) ≥100 kb in length and ran HumAS-HMMER (https://github.com/fedorrik/HumAS-HMMER_for_AnVIL) to identify alpha-satellite higher-order repeat (HOR) structures, thereby defining contigs containing complete centromeric structures.

### Various annotations on the DSA

Repeat element annotation was created using Rhodonite (10.5281/zenodo.6036498), a snakemake workflow combination of RepeatMasker v4.1.5 (http://www.repeatmasker.org), Tandem Repeats Finder (TRF^49^) v4.09 and DupMasker^50^, with a whole set of diploid DSA fasta as an input. Specifically for annotations marked by “Satellite” class by the RepeatMasker, we applied the following re-assignment of repeat name: (CATTC)n and (GAATG)n to HSat3, SAR to HSat1A and HSATI to HSAT1B. RepeatMasker annotation was then used for soft-masking of the DSA. Subsequently we defined CpG islands regions in DSA based on the soft-masked fasta file using cpg_lh in UCSC Kent utility. In order to identify the genic region in the diploid DSA, we implemented LiftOff^51^ v1.6.3 to project GENCODE^52^ V47 annotation. We first separated out the diploid DSA into the different haplotypes and ran LiftOff (*-copies -sc 0.95 -mm2_option=“-a -- end-bonus 5 --eqx -N 50 -p 0.5” -polish -cds -exclude_partial*) on an individual haplotype, respectively. We merged the resulting projected annotation files together subsequently to obtain a full annotation for the diploid assembly.

### Donor-specific graph (DSG) construction and projection of reads and variants between assemblies

Performing inter-haplotype comparison of SNV calls and regulatory features is challenging as each haplotype is defined by a unique coordinate system. To perform such comparisons, we first constructed a donor-specific pangenome graph (DSG) using Minigraph-Cactus^53^, without splitting by chromosome, clipping large unaligned regions, or filtering rare variants. This graph contains four assemblies, encoding the homology relationships between the two haplotypes of the COLO829BL cell line, as well as T2T-CHM13 and GRCh38. We then built a toolkit that enables us to move both genomic intervals (e.g. SNV calls, FIRE peaks) and entire mapped reads from one set of assembly coordinates to another. This enables us to move reads and features between the two haplotypes of COLO829BL or to move all features into a common coordinate space (e.g. GRCh38). This step makes use of the variation graph toolkit^54^ (vg *inject* and vg *surject* commands), along with custom pre- and post-processing steps. We use this framework to filter erroneous sSNVs arising from mismapping (see section "Diploid DSA-based somatic SNVs calling"), to bolster power when calling regulatory elements (see section "FIRE calling using the donor specific assembly" ), and to detect haplotype-distinct regulatory elements. To identify assembly segments unique to the DSA that are not present in GRCh38, we used halLiftover^55^ to detect regions with no alignment to GRCh38 in the DSG graph space.

### Alignment and post-processing of reads

Fiber-seq data were aligned against diploid COLO829BL DSA using pbmm2 (*--preset HiFi*) version 1.14.99, a wrapper for minimap2. Aligning the reads onto the diploid genome assembly would result in alignment records with zero mapping quality score (MAPQ) values, despite the reads originating from the same genome. For example, reads originating from either haplotype in homozygous stretches that exceed read length would be randomly assigned to one of the two haplotypes by the minimap2 alignment algorithm and MAPQ of those records will be set to zero. This may impact downstream analyses due to the fact that many genomics algorithms discard alignment below a certain threshold such as in variant calling, resulting in loss of genomic information. To avoid this issue, we arbitrarily set the MAPQ of all alignment reads to 60 using pysam (https://github.com/pysam-developers/pysam) and stored the original MAPQ reported by the aligner to a custom tag mq. CpG methylation data from fiber-seq data were extracted using pb-CpG-tools v3.0.0 (https://github.com/PacificBiosciences/pb-CpG-tools) *aligned_bam_to_cpg_score* command with default parameters. When running NucFlag using COLO829BL Fiber-seq data, reads with at least one of these following bitwise flags (unmapped (0×4), not primary alignment (0×100) and supplementary alignment (0×800)) were discarded using samtools (*view -F 2308*). Standard ONT and UL-ONT reads were aligned onto the DSA using minimap2 version 2.28-r1209 with the following parameters: *-ax map-ont -*y. Illumina short-reads and Element Aviti short-read sequencing data were aligned using bwa^56^ version 0.7.18 (bwa *mem*) and duplicated reads were marked by samblaster^57^ subsequently. Similar to Fiber-seq, MAPQ values in standard ONT, UL-ONT, Illumina and Element data were resetted.

### Identifying CDRs and quantifying their change

We merge alpha-satellite arrays within 10 kb of one another using bedtools^58^ and subsequently filter out all arrays shorter than 100 kb in length. We then filtered respective R9 UL-ONT files for alpha-satellite spanning reads using *samtools view -b -F 2308*. We generated bulk CpG methylation tracks across all reads using the fibertools^59^ *pileup* function with flags *–ml 225 –cpg*. To identify CpG methylation events along single molecules we used fibertools *extract* with flags *-r –ml 225 –cpg*. To identify “bulk CDRs” within Alpha-Satellite arrays, we binned CpG methylation bigWig values at 1 kb, computing per-alpha satellite stretch z-scores (using the specific array’s own mean and SD), and labeled bins with z-score < -1.5 as putative CDRs. Short runs (≤5 kb) that were flanked by the opposite state were smoothed, and only contiguous low-methylation segments at least 50 kb away from the interval edges were retained and adjacent segments were merged. The resulting segments were written as BED3 records for downstream analysis. To identify putative CDRs along single-molecules, we utilized a similar approach as the one used for calling bulk CDRs. Briefly, within each alpha-satellite array we segment each individual molecule into 1kb bins, and then calculate a per–read, per-bin z-score. Bins with Z less than -1.5 are then smoothed and merged with adjacent/overlapping segments, and output as a bed12 interval per molecule. Outputs were then subsequently visualized with the UCSC genome browser^60^ or IGV^61^. To identify the “center of mass” across CDRs for each chromosome we calculated the weighted average of each “bulk CDR” midpoint where the weight corresponds to the size of each CDR.

### Determining somatic haploid copy number (hCN) in COLO829

We derived the genome-wide somatic haploid copy number (hCN) state of COLO829 by leveraging the Fiber-seq coverage profiles from COLO829BL and COLO829 mapped onto the DSA and by comparing them based on the ground that COLO829BL exhibited normal karyotype. First, the DSA was segmented into 100 kb windows, and we excluded potentially misassembled regions. We then used mosdepth^62^ to calculate median coverage for each window (*--fast-mode --use-median --by*) for COLO829BL and COLO829 Passage B, respectively. For each window, we calculated the log2 ratio between COLO829BL and COLO829 Passage B while accounting for genome-wide coverage by normalizing each with normalization constant. To identify the constant, we inspected base-level coverage distribution for each input using mosdepth (Fig. S1b). As expected, COLO829BL Fiber-seq data showed a unimodal distribution with the mode at 176×. In contrast, COLO829 Passage B Fiber-seq data exhibited a bimodal distribution for COLO829 Passage B Fiber-seq data, with primary mode at 123×. Based on prior observations of the COLO829 karyotype, which has undergone multiple whole-genome duplication events, we reasoned that the mode is indicative of the reads in the duplicated segments, and the second peak indicates haploid (1n) coverage. By using haploid (1n) coverage as normalization constant controlling for genome-wide coverage, we were able to estimate the genome-wide log_2_hCN ratio as follows:

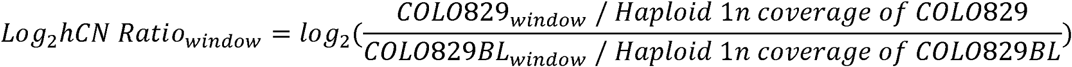

Next, after capping log_2_hCN ratio values below –1.5 at –1.5, we employed a circular binary segmentation (CBS^63^) algorithm implemented in the DNAcopy^64^ package to identify changes in copy number states across COLO829 genome with the following step: (1) *smoothed_log2hCN <-smooth.CNA(log2hCN, smooth.region=10, outlier.SD.scale=4, smooth.SD.scale=25, trim=0.025)*; (2) *cbs_result <-segment(smoothed_log2hCN, min.width = 5, nperm = 10000, alpha = 0.0001)*. Next, we assigned hCN to each segment based on the median value of the merged segments. Segments were classified using breakpoints calculated as log_2_((n+0.5)), where n represents the haploid copy number state (0 through 4). This approach accounts for the expected log2 ratio for each haploid copy number relative to a haploid baseline. Specifically, segments with median log_2_hCN ratios≤–0.415 (log2(0.5)) were classified as haploid deletions (DEL); ratios>–0.415 and ≤0.585 (log_2_(1.5)) as single-copy (hCN=1); ratios>0.585 and ≤1.322 (log_2_(2.5)) as single-copy gain (hCN=2); ratios>1.322 and ≤1.848 (log_2_(3.5)) as two-copy gain (hCN=3); ratios>1.848 and ≤2.263 (log_2_(4.5)) as three-copy gain (hCN=4); and ratios >2.263 as high-level amplification (hCN>4). These breakpoints represent the midpoints between integer copy number states, minimizing misclassification due to technical noise or heterogeneity. Then, the proportion of the genome altered by somatic copy number alteration was then calculated using the rustybam^65^ *bed-length* command on a 100kb window bed file that specified the somatic hCN state for each segment. Furthermore, to determine cross-haplotype hCN information, we leveraged the DSG to move reads assigned to one haplotype to the corresponding region on the other haplotype. We then applied an identical hCN discovery approach using the coverage profile of these haplotype-switched reads.

To identify the chromosomal-level genomic rearrangement events and pinpoint their locations, we manually curated the changepoints in the contigs and examined the supplementary alignment records for UL-ONT, standard ONT and Fiber-seq alignments for COLO829. This enabled us to determine the base-resolution locations of the rearrangement events such as t(1;3)(q12;p21). Furthermore, we checked whether or not the translocation events of i(4)(q10) and t(14;16)(p10;p10), each of which involves a breakpoint in an alpha-satellite HOR element could occur by chance alone. Statistical significance was assessed using a Monte Carlo permutation test with 100,000 iterations. The permutation test p-value represents the probability of observing ≥2 overlaps by chance alone.

### Diploid DSA-based somatic SNVs calling

To establish diploid DSA-based somatic single nucleotide variants (sSNVs) set in COLO829 using Fiber-seq data, we first called SNVs using DeepVariant^66^ v1.6.1 separately on COLO829BL Fiber-seq and two Fiber-seq data from COLO829 biological replicates. We devised multiple filtering schemes to ensure that we identify the fixed cancer variants of high quality. First, we filtered out SNV calls that are overlapped with potentially misassembled regions, we retained only those that are present in both COLO829 replicates but not present in the COLO829BL. Additionally, we applied a pileup-based filtering scheme such that we allow only single alternate allele to be present in the COLO829BL sample in the SNV records shared by two replicates (Figure S3A). We discovered that once we push the threshold, the spurious mutational signatures arise starting from alternate read depths of two (Figure S3B), which means that we are effectively removing potential artefacts arising from sequencing error and those arising from prolonged culture of the cell line. Next, we removed SNV calls overlapped with deleted segments determined by haploid copy number analysis (Figure S3A), which are resulting from genomic stretches homozygous from the corresponding intact haplotype. We then set out to resolve sSNVs which are called in both haplotypes at homologous regions. As identical sSNVs are highly unlikely, these duplicated SNVs likely arise from highly homozygosity between haplotypes, or segmental duplications within a haplotype, resulting in mismapping of reads from one haplotype to the other. To identify these sSNV pairs, we leveraged the DSG (Figure S3A), moving sSNV calls to their non-native haplotype and looking for overlapping pairs. To determine which of these is likely the ’true’ variant, we used a multi-step filtering approach. We first applied bcftools *mpileup*^67^ on UL-ONT data to calculate the fraction of reads supporting the sSNV and reference base in each haplotype. We then performed a Fisher’s exact test on these values and assigned an sSNV to one haplotype for records with a *p*-value<0.01. We re-processed the remaining records in the same fashion, but now using ONT data as input to bcftools *mpileup*. SNV records that remained ambiguous were flagged and excluded from certain analyses but retained if not stated. Finally, we applied an SNV density filtration method (Figure S3A). We leveraged the fact that false positive sSNV records were predominantly found to be derived from alignment artifacts localized near structural variations, particularly deletions during the visual inspection of sSNVs using karyoploteR^68^. Hence, we segmented the genome into 1 kb window with a 500 bp sliding window approach and intersected the remaining sSNV records with these windows. We selected windows containing 3 or more sSNVs and merged them. sSNVs were discarded from windows if three or more sSNVs in that window each had a VAF below 0.75, or if the median read coverage for sSNVs in the window was below the threshold defined as 61.5–3×σ, where σ is the standard deviation of a Poisson distribution with μ=61.5, representing the haploid coverage of COLO829 Fiber-seq data (hCN=1) on which our SNV calling was based. Subsequently, validation of sSNVs using orthogonal sequencing data was performed using bcftools *mpileup*.

### Mutational spectrum analysis for sSNV and sDNV

Comprehensive mutational spectrum analysis was performed to characterize the patterns of somatic SNVs (sSNVs) and dinucleotide variants (sDNVs) across our dataset. While multiple established software packages exist for variant call format (VCF) processing and mutational signature analysis, we developed a custom Snakemake workflow (https://github.com/ryansohny/VCF2SPECTRUM) that integrates existing bioinformatics softwares with custom codes to ensure reproducibility and scalability across VCF data based on diploid genome alignment.

### Generating SBS-96 and DBS-78 mutational matrix

The construction of single base substitution (SBS) matrices required systematic extraction and categorization of mutations within their sequence contexts. The process involved two sequential mutyper^69^ commands: mutyper *variants* to extract trinucleotide contexts (*k*=3) around sSNVs from the DSA, followed by mutyper *spectra* with the *--population* parameter to generate mutation spectra. Then, raw mutyper output was converted to SBS-96 format compatible with SigProfilerAssignment^70^. Identification of DBS began by extracting all sSNVs separated by exactly 1 base pair, focusing on clusters containing even numbers of variants to identify potential sDNV events. For variants with homozygous genotypes (1/1;GT field), direct assignment as DBS was performed when adjacent SNVs were identified. However, heterozygous GT combinations required additional validation: when two adjacent heterozygous GT were detected, WhatsHap^71^ v2.8 was employed for read-backed phasing to confirm that both substitutions occurred on the same DNA molecule, ensuring the identification of true sDNVs rather than independent sSNVs on different alleles. We then performed manual visual inspection of the reads using IGV^61^ to verify the phasing and confirm the presence of genuine sDNVs.

### Normalization of SBS-96 matrix by expected K-mer frequencies

To account for differences in trinucleotide k-mer (3-mer) frequencies (Figure S5A) and mitigate the resulting sequence context bias, where certain mutation types appear more frequently simply due to greater availability of their target sequences, SBS-96 matrix were normalized before assignment of known COSMIC SBS signatures^72^. 3-mer frequency for each selected region of the genome was identified through Jellyfish^73^ (*count -m 3 --canonical*). Normalization factors were calculated as the ratio of genome-wide to region-specific 3-mer frequencies, then applied to raw mutation counts. To preserve the total mutational count in the original matrix, a global scaling factor was calculated and applied to all normalized counts, with final values rounded to integers. To compute region-specific predicted mutability, we first derived 3-mer enrichment vectors by normalizing each 3-mer frequency matrix against the 3-mer distribution in non-repetitive regions. We then calculated the adjusted SBS-96 mutational spectrum fraction for each specified region by multiplying the observed fraction of each SBS-96 mutation type by its region-specific 3-mer enrichment factor. The sum of adjusted fractions across all 96 substitution types yields the predicted mutability, quantifying the expected relative mutation rate after correcting for underlying sequence composition bias.

### Mutational signature assignment

Mutational signature assignment was performed to decompose the 3-mer-corrected mutational spectra (SBS-96 only) into known signatures catalogued in the COSMIC database^72^ v3.4 using SigProfilerAssignment^70^ *cosmic_fit()* function, which uses non-negative least squares regression. To avoid overfitting, reconstructing the SBS-96 matrix was done using signatures previously known to be associated with melanoma^22^: SBS1, SBS2, SBS5, SBS7a/b/c/d, SBS13, SBS17a/b, SBS38, SBS40 and SBS97, unless stated otherwise.

### Aggregate Mutation Spectrum Distance (AMSD) analysis

To test whether 3-mer SBS mutation spectra (i.e., SBS-96) differed between genomic contexts (e.g., non-repetitive genome vs. satellite elements), we applied the aggregate mutation spectrum distance (AMSD; Figure S5B) method^74^. Briefly, AMSD compares the cosine distance between two observed spectra with the distribution of cosine distances obtained from random re-sampling of mutations from the same spectra, treating each mutation as independent. The resulting *p*-value reflects the fraction of random re-samplings that produced a greater cosine distance than observed. Mutation counts were corrected to the expected 3-mer distribution of the full genome prior to AMSD analysis, same as described above for mutational spectrum analysis for sSNV. After correction, we ran 100,000 AMSD permutations per comparison, giving a minimum possible p-value of 1×10⁻LJ.

### Detecting somatic restructuring of telomeres

We identified telomeric regions in the DSA by using the seqtk *telo* tool. We then use fibertools *center* with the *-w* flag and the seqtk *telo* bed file to place single-molecules in a coordinate space relative to the telomere/sub-telomere boundary. We subset our analysis to telomeres in the DSA with a clearly unique reference sequence. We quantified the relative length of all telomeres by measuring the number of bases that appear in each telomeric read after the subtelomere-boundary alignment position. We applied a regular expression to identify canonical CCCTAA repeats. We then deduced the strand of origin of the read by counting the number of CCCTAA or TTAGGG instances, and the motif with more instances was assigned as the canonical motif. Utilizing the positions of the canonical motifs we identified stretches of bases that were not part of a CCCTAA repeat (TVRs). As a first pass for identifying telomeric restructuring events we build a consensus TVR structure for each telomere in COLO829BL and COLO829 that represents the percent of bases at each position that is a TVR. We then apply a 25 bp smoothing function and identify TVR “peaks” in both cell lines. We then search for the first peak that is present in COLO829BL but not COLO829 as the distal “boundary” delineating the minimum amount “chewed back”. The first shared peak between the two cell lines is the sub-telomeric proximal limit for the maximum amount “chewed back” during telomeric restructuring. We performed this in both ONT (R10) and PacBio HiFi reads and a telomere was only deemed restructured if the restructuring event was detected by both technologies. This first pass identified 12 putative telomeres which we then visually inspected and deemed that only 10 had undergone a clear TVR restructuring event.

### FIRE calling using the donor specific assembly

Calling FIRE peaks using reads mapped to a diploid donor-specific genome preserves haplotype-specific variants, but effectively halves the power to detect peaks relative to using a haploid reference. To resolve this, we use the DSG to lift mapped reads between genome assemblies using vg inject and vg surject (version 1.68). Each read is lifted to its non-native haplotype, resulting in a merged output file containing both the native and lifted coordinates for each read. The final output thus contains "diploid" coverage on each haplotype and serves as input for FIRE peak calling. Additionally, this approach makes it straightforward to detect haplotype-selective FIRE peaks, based on relative haplotype coverages at each peak. FIRE peaks called in DSA space can themselves be lifted between the assemblies in the DSG using the approach above, enabling us to lift peaks between haplotypes to compare their positions, or to a common reference assembly (GRCh38).

## Data and code availability

The SMaHT Data used in this study can be accessed from the SMaHT Data Portal at https://data.smaht.org/. For the SMaHT donor tissue, all protected (e.g., sequencing data, germline variant calls, and complete donor metadata for the SMaHT Production donors) and open (e.g., gene expression quantification, tables, etc.) access data are available in dbGaP (phs004193 for the SMaHT Benchmarking data; phs004104 for the SMaHT Production data. The original code to reproduce all of the data processing and analysis in this study can be found at https://github.com/StergachisLab/T2T-COLO829BLT-Manuscript. The results and annotations are available as trackHubs on https://genome.ucsc.edu/cgi-bin/hgTracks?db=hub_6753970_DSA_COLO829BL_v3.0.0.

## Acknowledgements

We thank members of the SMaHT consortia as well as Steve Henikoff for their helpful feedback on this manuscript. This research is supported by the NIH Common Fund, through the Office of Strategic Coordination/Office of the NIH Director under awards U24 MH133204, U24 NS132103, UG3 NS132024, UG3 NS132061, UG3 NS132084, UG3 NS132105, UG3 NS132127, UG3 NS132128, UG3 NS132132, UG3 NS132134, UG3 NS132135, UG3 NS132136, UG3 NS132138, UG3 NS132139, UG3 NS132144, UG3 NS132146, UM1 DA058219, UM1 DA058220, UM1 DA058229, UM1 DA058230, UM1 DA058235, and UM1 DA058236. A.B.S. holds a Career Award for Medical Scientists from the Burroughs Wellcome Fund and is a Pew Biomedical Scholar. This study was supported by National Institutes of Health (NIH) grants 1DP5OD029630, and 1U01HG013744 to A.B.S.. M.R.V. and S.C.B. were supported by a training grant (T32) from the NIH (2T32GM007454-46). M.R.V. was also supported by a Pathway to Independence award from the National Institute of General Medical Sciences (1K99GM155552-01). D.D. was supported by a training grant (T32) from the NIH (T32GM141828).

## Supplemental Figure Legends

**Figure S1. Building near-T2T diploid DSA of COLO829BL and its quality metrics.**

**A.** Detailed diagram of constructing COLO829BL DSA and identifying somatic genomic events leveraging the assembly.

**B.** Comparison of contiguity of the genome assembly using NG50 statistics.

**C.** Quantitative measurement of assembly completeness across the DSA haplotypes, GRCh38 and T2T-CHM13.

**Figure S2. Chromosomal rearrangements and haploid copy number alterations revealed by DSA-based analysis.**

**A.** Coverage histogram of COLO829BL and COLO829 (Passage B) Fiber-seq data across the diploid DSA. The coverage for 10 million bases was randomly drawn for each sample with sampling weighted by the length of the interval and excluding zero coverage regions.

**B.** An example of identifying chromosomal-level rearrangements between chromosome 1 and chromosome 3 using log2hCN profiles across the DSA contigs.

**C.** Schematic reconstruction of complex genomic rearrangements between chromosome 1 and 3 identified by the DSA-based analysis.

**D.** Log2hCN profiles of q-arm of two chromosome 4 haplotypes forming isochromosome.

**E** and **F.** Log2hCN profiles of chromosome 14 and 16, together with triplication and duplication events, respectively. The color of the ideograms on top of log2hCN ratio plot represents the haplotype of origin (haplotype 1:blue; haplotype 2:red). Each green dot represents a 100kb window “marker” with a red line indicating the median of each segment.

**Figure S3. Comprehensive somatic single nucleotide variant discovery and validation using the DSA.**

**A.** Sequential methods for identifying and refining sSNVs in COLO829 using the DSA. We integrated somatic haploid copy number data, a donor-specific graph (DSG), and density-based filtering to obtain a final set of 44,795 sSNVs and 828 sDNVs (see Methods).

**B.** Heatmap showing the relationship between potential false-positive sSNVs and mutational signature composition. Each column represents the number of COLO829BL reads matching the tumor alleles, indicating likely false positives variants. Rows represent COSMIC single base substitution (SBS-96) mutational signatures (For this particular analysis, mutational spectrum normalization using 3-mer frequency (see Methods) was not applied and whole COSMIC SBS-96 signatures were used to reconstruct each mutational spectrum).

**C.** Orthogonal validation rate of COLO829 sSNVs stratified by read support.

**D.** IGV view of sSNVs present in chromosome 1 at the translocation breakpoints between chromosomes 1 and 3. Red arrows indicate sSNVs with ∼0.5 VAF exclusive to the translocated fragment; green arrows mark sSNVs unique to the intact chromosome 1; blue arrows denote sSNVs present in both the intact chromosome 1 and the translocated segment.

**E. h**VAF histogram of sSNVs stratified by involvement in translocation t(1;3). Lime green: sSNVs on chromosome 1 p-arm (not involved in translocation); salmon: sSNVs on chromosome 3 p-arm (involved in t(1;3) translocation); magenta: sSNVs on chromosome 1 q-arm (present on four copies, two of which are translocated to chromosome 3).

**Figure S4. Haploid variant allele fraction distribution pattern across genomic contexts.**

**A.** hVAF distribution of sSNVs across **A.** different chromosomal contigs in the COLO829BL DSA.

**B.** different repeat classes and **C.** different chromosomal contigs separated by hCN states.

**Figure S5. Mutational spectrum analysis across various genomic contexts.**

**A.** 3-mer fraction across non-repetitive portions of the DSA and regions with different repeat elements.

**B.** AMSD analysis results which compared mutational spectra between non-repetitive regions and different repetitive elements.

**C.** Relationship between predicted mutability and observed sSNV mutation rate for relative to non-repeat elements across different repeat element classes.

**Figure S6. Pattern of somatic SNVs in alpha-satellite regions of COLO829 centromeres**

**A.** IGV tracks showing sSNV distribution in CDR and non-CDR alpha-satellite regions for chromosomes 17 and 22.

**B.** 3-mer fraction across CDR and Non-CDR alpha-satellite is largely identical with a cosine similarity of 0.99945.

**C.** Predicted mutability vs. observed sSNV mutation rate for alpha-satellite repeats (CDR and non-CDR combined) relative to non-repeat elements.

**Figure S7. Genome browser view of somatic rewiring of CDRs in COLO829.**

**A-F** Genomic loci displaying CDR restructuring. For each locus, in order from top to bottom, are alpha-satellite location, mCpG %, FIRE Accessibility %, Bulk CDR calls, FIRE peaks, and sing-molecule CDR calls. Top-most tracks represent COLO829BL, bottom tracks represent COLO829 (Passage B)

**Figure S8. Single-molecule-resolution architecture of telomeric restructuring events in COLO829.**

**A.** Restructured telomeric loci detected with PacBio Sequencing. Every individual bar represents a single-molecule sequencing with PacBio HiFi sequencing. Blue boxes represent canonical telomeric motifs (TTAGGG_N_), and red boxes indicate bases that cannot be classified within the canonical motif. The left most position for each individual read represents the boundary between sub-telomere and telomere, identified with *seqtk telo*. For every locus (left), COLO829BL is displayed on top, COLO829 displayed below.

**Figure S9. Relationship between somatic alterations and epigenome rewiring.**

**A.** Identifying Fiber-seq-inferred regulatory elements (FIREs^10^) in the genome using Fiber-seq and fibertools.

**B.** Associating FIREs in two corresponding haplotypes by leveraging the donor-specific assembly graph (DSG) constructed with T2T-CHM13, GRCh38 and COLO829BL diploid DSA.

**C.** FIRE calls on individual reads at sites with haplotype-selective chromatin accessibility (HSCA) and a somatic variant.

**D.** Difference in accessibility between the two haplotypes for sites with HCSA, both with and without intersecting somatic variants.

**E.** Comparison of chromatin accessibility between haplotypes where one of the haplotypes has been amplified in COLO829 compared to the original COLO829BL copy number seen in the normal haplotype.

**F.** The fraction of haplotype-selective Fiber-seq peaks without somatic variants or heterozygous germline variants (dark red) or other HSCA peaks (red) located within a specific distance (Mbp) from a change in copy number along the COLO829 genome and their difference from the cumulative distribution of all other FIRE peaks (see Figure 6D).

